# The haplolethal gene *wupA* of Drosophila exhibits potential as a target for an X-poisoning gene drive

**DOI:** 10.1101/2023.06.23.546292

**Authors:** Clancy D. Lawler, Ana Karla Parra Nuñez, Natalia Hernandes, Soumitra Bhide, Isabelle Lohrey, Simon Baxter, Charles Robin

## Abstract

A synthetic gene drive that targets haplolethal genes on the X-chromosome can skew the sex ratio towards males. Like an ‘X-shredder’ it does not involve ‘homing’ and that has advantages including the reduction of gene drive resistance allele formation. We examine this ‘X-poisoning’ strategy by targeting four of the 11 known X-linked haplolethal/haplosterile genes of *Drosophila melanogaster* with CRISPR/Cas9. We find that targeting the *wupA* gene during spermatogenesis skews the sex ratio so fewer than 14% of progeny are daughters. That is unless we cross the mutagenic males to X^^^XY female flies that bear attached-X chromosomes, which reverses the inheritance of the poisoned X chromosome so that sons inherit it from their father; in which case only 2% of the progeny are sons. These sex ratio biases suggests that most of the CRISPR/Cas9 mutants we induced in the *wupA* gene are haplolethal but some are recessive lethal. The males generating *wupA* mutants do not suffer from reduced fertility rather the haplolethal mutants arrest development in the late stages of embryogenesis well after fertilized eggs have been laid. This provides a distinct advantage over genetic manipulation strategies involving sterility which can be countered by the remating of females. We also find that *wupA* mutants that destroy the nuclear localization signal of shorter isoforms are not haplolethal as long as the open reading frame remains intact. Like *D. melanogaster wupA* orthologs of *D. suzukii* and *Anopheles* mosquitos are found on X chromosomes making *wupA* a viable X-poisoning target in multiple species.

## Introduction

For many decades it has been envisioned that insect populations that vector diseases or that are pests of crops could be controlled using genetic methods (Serebrovsky 1969; Knipling 1955; Curtis 1968; Whitten 1985; Burt 2003). Indeed, Sterile Insect Techniques have successfully controlled fly species (Bushland, Lindquist, and Knipling 1955; Hendrichs, Franz, and Rendon 1995; Scott *et al*. 2017) and field trials relying on self-limiting transgenes are currently being deployed and assessed (Yao *et al*. 2022; Waltz 2021). Recently, the prospect of synthetic gene drives, a distinct class of population manipulation, where transgenes can be engineered to spread through otherwise natural populations of insect pests has received much attention (Alphey *et al*. 2020; Hay, Oberhofer, and Guo 2021; Champer, Buchman, and Akbari 2016). The effectiveness of various gene drive designs, many of which use CRISPR/Cas9 machinery, have been assessed in laboratory populations (Chen *et al*. 2007; N. Windbichler *et al*. 2007; Akbari *et al*. 2014; Kyrou *et al*. 2018; Oberhofer, Ivy, and Hay 2019; Webster, Vella, and Scott 2020; Yang *et al*. 2022). Many of these require ‘homing’ where double-stranded DNA breaks must be repaired by homologous recombination (HR) so that the transgenic selfish elements are copied and thereby spread. A drawback of such homing designs has been that some double stranded breaks can be repaired using the non-homologous end-joining (NHEJ) pathway. In these instances not only does homing fail but mutations that are resistant to subsequent CRISPR/Cas9 cleavages can be introduced into the population (Nikolai Windbichler *et al*. 2011; Gratz *et al*. 2014; Champer *et al*. 2017; 2018; Hammond *et al*. 2017).

One genetic control design that does not require homing is referred to as the X-shredder (Galizi *et al*. 2014). It targets highly repeated sequences on the X chromosome during spermatogenesis such that X-bearing sperm are inviable, and the drive skews the sex ratio towards males. If the X-shredding transgenes could be placed on the Y chromosome, then an inheritance can be established such that fathers only have sons. Originally, the X-shredder was attempted in lab populations of *Anopheles gambiae* mosquito using the endonuclease *PpoI*, but subsequent implementations in *A. gambiae* mosquitos, *Drosophila melanogaster and Ceratitis capitata* fruit flies have deployed CRISPR/Cas9 machinery (Galizi et al. 2014; Fasulo et al. 2020; Meccariello et al. 2021; Haber et al. 2023).

A related approach, dubbed ‘X-poisoning’, also targets loci on the X chromosome; not so the chromosome is physically destroyed but so that mutations are introduced such that progeny receiving the X chromosome will not be viable (Burt and Deredec 2018; Fasulo et al. 2020; Haber et al. 2023). This strategy uses CRISPR/Cas9 to target X-linked haplolethal genes. Although haplolethal genes are typically thought of as genes where both alleles of a diploid organism need to be functional, X-linked haplolethal genes can occur in many dipterans because their dosage compensation mechanism elevates the X chromosome-encoded output in males (that is in contrast to eutherian mammals where there is silencing of X loci in females and therefore haplolethals on the X would be incompatible with maleness) (Rose *et al*. 2016; Cook *et al*. 2012). There may be a few potential advantages of X-poisoning over X-shredding. Firstly, X-poisoning does not rely on natural repeats occurring exclusively on the X chromosome and theoretically this may increase its applicability to some species. Secondly, CRISPR cutting in one or a few sites on the X chromosome might occur more readily than cutting the X chromosome at an overwhelming number of places, as is required for X-shredding. Thirdly, X-shredding has been shown to be ameliorated by the NHEJ pathway thus potentially limiting its utility (Fasulo *et al*. 2020). Fourthly, it may be that post-zygotic effects of X-poisoning may change the spread dynamics so it has a higher threshold than X-shredding and that may lead to more controllable population manipulation.

Here we describe the targeting of haplolethal genes on the X chromosome of *Drosophila melanogaster* with CRISPR/Cas9. The loci differ from the two targeted in the X-poisoning study of Fasulo *et al*. (2020) as we examine three other ribosomal protein encoding genes and the haplolethal gene *Wings up A (wupA)*. In contrast to the ribosomal protein genes which might be required for the viability of most cell types including those in spermatogenesis, *wupA*, encodes a troponin that is expressed in muscles from mid embryogenesis. We therefore sought experimental affirmation that these genes are indeed haplolethal genes and whether they could be valuable targets for X-poisoning gene drives.

## Materials and Methods

### gRNA design and cloning

A combination of ChopChop v3 (Labun *et al*. 2019) and CRISPR Optimal Target Finder (Gratz *et al*. 2014) was used to identify the gRNA target sites with the highest efficiency scores with a low likelihood of off-target activity. All gRNA sequences were checked against the DGRP reference panel using FlyBase’s JBrowse (Larkin *et al*. 2021) to avoid standing variation at the target site that may give natural resistance.

The design protocol of Port and Bullock (2016) was used to clone gRNA sequences into the pCFD5_w plasmid (Addgene #112645). Three separate plasmids were generated that targeted haplolethal genes on the X chromosome; pRpS19a_RpL35, pRpL36_RpL35, and p*wupA*. pRpS19a_RpL35 encoded four gRNAs, two targeting *RpS19a* (*RpS19*_RNA1: GAAGGATATTGACCAGCACG, *RpS19*_RNA2: AACCGACTCCAGCGGGACTG) and two targeting *RpL35* (*RpL35*_gRNA1: GTGCTCCGAGCTGAGGATCA, *RpL35*_gRNA2: CACAATGTAGACGCGAGCGA). pRpL36_RpL35 encodes four gRNA, two targeting *RpL36* (*RpL36*_gRNA1: GCTGGCTATTGGCCTGAACA, *RpL36*_gRNA2: CATGCGCGACTTGGTCCGCG) and two targeting *RpL35* (as aforementioned). Each of the three sgRNA’s used to target *wupA* (*wupA*_gRNA1: ACCAAAAACACAAATCAAAA, *wupA*_gRNA2: TGAGGTGCGCAAGCGCATGG, *wupA*_gRNA3: CGCATCATCGAAGAACGTTG) would affect all 13 alternate transcripts annotated in Flybase. The PAM site for *wupA*_sgRNA1 corresponds to the most common start codon and is in the 5’UTR of transcripts using the alternate start codon, the PAM site for *wupA*_sgRNA2 corresponds to the alternate *wupA* start methionine which is a codon used by in all transcripts, and that for *wupA*_sgRNA3 is in a coding exon common to all transcripts (Figure 1). All primers used and sgRNA sequences are tabulated in Table S1.

**Figure 1:**
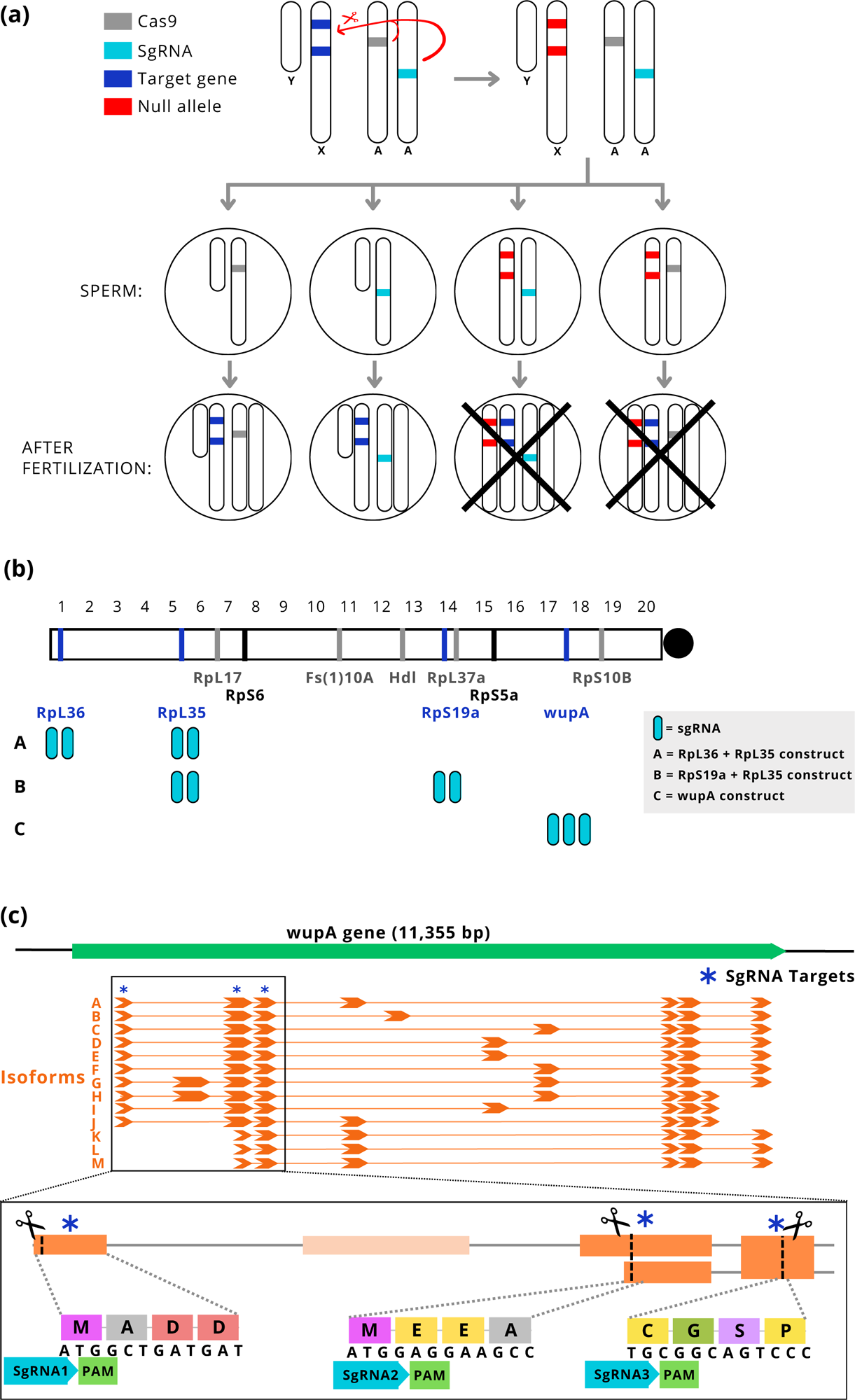
Targeting Haplolethal genes on the X chromosome. (a) Premeiotic cells expressing CRISPR/Cas9 (from the autosomes) that effectively target haplolethal loci on the X-chromosome (blue to red) will result in male only offspring. If the machinery were put on the Y-chromosome then inheritance of the elements will be limited to the male germ line and a gene drive would be created. (b) Eleven haplolethal/haplosteriles were reported by Cook *et al*. (2012), two (RpS6 and RpS5a) were targeted by Fasulo *et al*. 2020 and the location of the four loci targeted in this manuscript, and the number of sgRNA used (rounded blue rectangles) are shown. (c) The *wupA* gene encodes for thirteen transcript isoforms. SgRNA1-3 target three conserved regions of the gene’s coding sequence.

### Microinjections

250-500 ng/uL of the pRpS19a+RpL35 and p*wupA* plasmids were microinjected into the 09w fly line while the pRpL36+RpL35 was injected into the 10w fly line. These two fly lines were obtained from Trent Perry, Bio21 Unimelb who had generated them using the Bloomington stocks #24749 and #25709 (y[1] v[1] P{y[+t7.7]=nos-phiC31\int.NLS}X; P{y[+t7.7]=CaryP} attP40) and #25710 (P{y[+t7.7]=nos-phiC31\int.NLS}X, y[1] sc[1] v[1] sev[21]; P{y[+t7.7]=CaryP}attP2) to generate second chromosome landing site flies (09W) and third chromosome landing site flies (10W) in a *white* null background. Insertion events were validated by sequencing and single integration lines were maintained.

### Assessing sex ratio biasing capabilities of sgRNA constructs

To assess the sex ratio biasing rates of each of the sgRNA expressing constructs, we first generated male flies that expressed both Cas9 and sgRNAs in their germ cells. Individual females homozygous for each of the three sgRNA constructs (RpS19a+RpL35, RpL36+RpL35 and *wupA*) were crossed to males containing (i) Cas9 under the control of the *nanos* promoter (Either NIG CAS-0001; y^2^ cho^2^ v^1^; attP40{nos-Cas9}/CyO which carries a recessive lethal allele on the second chromosome and cannot be made homozygous) or (ii) a line we generated that has nos-Cas9 (from Addgene plasmid 62208 courtesy of Simon Bullock which has a single NLS and a 3’UTR from *nanos*) placed in the 09w second chromosome attP40 landing site (referred to herein as nos-Cas9(II)_NH) or (iii) a *vasa*-cas9 on the Y chromosome from BL91386 (Gamez et al. 2021). The paternal inheritance of Cas9 prevented maternal deposition and somatic activity of the drive (Champer *et al*. 2019) and is consistent with the ultimate aim of placing X-poisoning transgenes on the Y chromosome. Initially, single males heterozygous for Cas9 and one of the sgRNA constructs were crossed to three 09w 3–4-day old virgins (14 replicates) and allowed to mate and lay in a vial for seven days before being cleared and all offspring that emerged counted. Subsequent crosses (14 replicates) included single nos-Cas9/*wupA*-sgRNA males left to mate with single females for four hours, or alternatively crossed to 3-day-old virgin females with the genotype C(1)DX,*y*^1^,*f*^1^/Y (Bloomington Stock number 4559; 7 replicates), which is an ‘attached-X’ line that forces paternal X chromosome inheritance.

### Analysis of life stage associated with *wupA* lethality

Next we assessed the lifestage in which sex skewing occurred with *wupA* X-poisoning crosses. Single males with a GFP marked X-chromosome (y[1] M{RFP[3xP3.PB] GFP[E.3xP3]=vas-int.Dm}Zh-2A w[*]/Y; sg(*wupA*)/nos-Cas9 were crossed to three wildtype virgin females (DGRP line 897) and allowed to mate for 48 hours. The control cross was the same except there was no Cas9. All flies where then transferred to a vial with grape juice agar and allowed to lay eggs for 14 hours, after which the flies were cleared. At the time of clearing, the total number of eggs laid were counted. Twenty-four hours later the eggs were sexed and the female eggs (GFP) were separated from the male eggs (non-GFP). We then tracked how many of the female eggs successfully emerged into larvae, pupae, and adults. We performed 20 replicates for each cross.

### Competitive mating assays

We assessed mate choice by placing one *ebony* (dark body colour) homozygous female in a vial with one *ebony* homozygous male and one test male of the following genotypes: A) sg(*wupA*)/nos-Cas9, B) sg(*wupA*)/+, C) +/nos-Cas9, D) 09w. The flies were allowed to mate for 24 hours, after which the parents were cleared and the progeny was phenotyped for body colour once they reached adulthood. All flies were 2-3 day old virgins at the time of mating.

### Sequence characterization of *wupA* survivors

Sons of crosses between X-poisoning males and attached-X females had their DNA extracted and were characterized with Sanger sequencing using standard protocols. PCR primers were used to amplify the DNA around all sgRNA targeted sites of a single gene, these were purified with a spin column and Sanger sequenced by Macrogen (see Table S1 for sequencing primers).

Two hundred of the surviving female progeny from the cross 09w x sg(*wupA*)/nos-Cas9 were collected and pooled into batches of ten. Their DNA was extracted following a standard phenol:chloroform DNA extraction protocol. PCR reactions were then performed using the primers labelled “*wupA* sg# Illumina sequencing” in Table S1. Each set of primers amplifies a specific sg(*wupA*) target site. The PCR products of each female sample were pooled together and purified with Bioneer AccuPrep PCR/Gel Purification Kit. Samples were sent for MiSeq 300 sequencing with AGRF. The sequencing results were input into Crispresso2 for allele-specific quantification of each sg(*wupA*) target site (Clement et al. 2019).

## Statistical Analyses

All statistical analyses were undertaken in R (R Core team 2023). Sex ratio biasing in the *Drosophila* X-poisoning gene drive crosses were analysed using an unpaired two-tail t-test with a Bonferroni correction for multiple comparisons where applicable.

## Results

### X-poisoning targeting ribosomal protein genes

Four of the haplolethal genes described on the X chromosome of *Drosophila melanogaste*r (Cook *et al*. 2012) were targeted using three multi-sgRNA constructs and Cas9 expressed in the male germline via a *nanos* promoter (Figure 1). The expectation was that males carrying the Cas9 and sgRNA transgenes would have a reduced number of viable daughters relative to controls yet the same number of sons (Figure 2a). All the crosses targeting ribosomal protein encoding genes (RpS19a+RpL35, RpL36+RpL35) yielded significantly fewer offspring when compared to the controls (*p*<0.0001, t-test with Bonferroni correction; Figure 2b; Table 1). The crosses involving the RpS19a+RpL35 sgRNA construct followed expectations with significantly fewer daughters than sons (*p*<0.0001, t-test with Bonferroni correction) and no significant difference in the number of sons compared to the control. In contrast the RpL36+RpL35 crosses yielded significantly fewer sons *and* daughters compared to control (*p*<0.05 and *p*<0.01 respectively, t-test with Bonferroni correction) without a significant sex ratio bias (Table 1).

**Figure 2:**
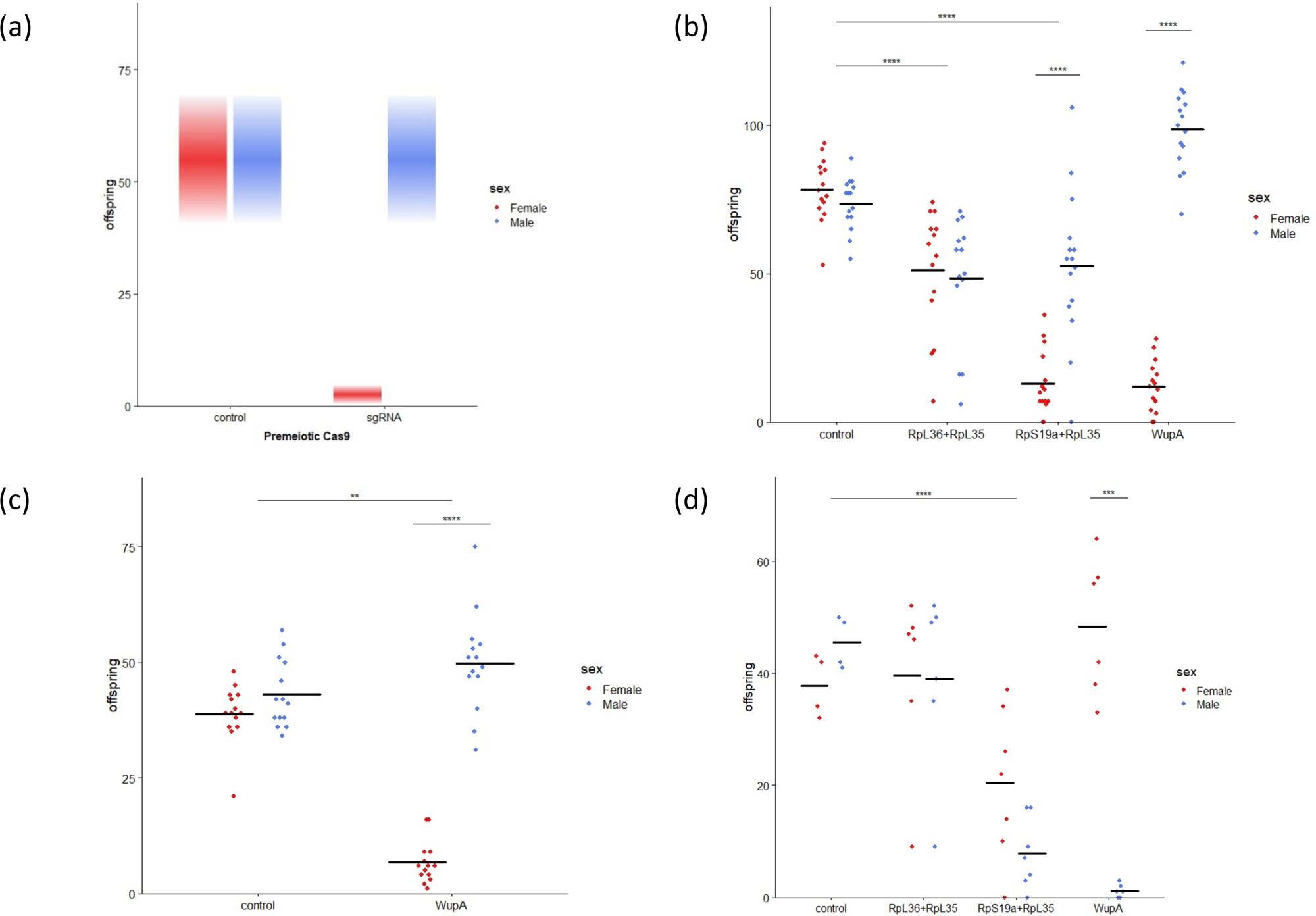
Sex biases with X-poisoning. (a) A symbolic representation of what is expected in our crosses. A skew against daughters (red) is expected when sgRNA’s target haplolethal loci on the X chromosome during gametogenesis. The number of sons is expected to remain unchanged as they would arise from sperm carrying the Y chromosome. (b) The observed amount of sex biasing skew ranged from nothing (RpL36+RpL35), to modest (RpL19a+RpL35), to promising (wupA). Note that the total number of offspring was also variously affected. (c) By excluding multiple mating the total number of offspring in the *wupA* cross was reduced and there was a strong skew towards males (d) CRISPR/Cas9 fathers crossed to attached-X mothers, reverse the inheritance of the sex chromosomes so that sons inherited the CRISPR modified X chromosome, and daughters inherited the Y chromosome. (* *P* < 0.05, ** *P* < 0.01, ** *P* < 0.001 **** P < 0.0001, unpaired two-tail t-test with a Bonferroni correction for multiple comparisons).

**Table 1:** Sex skews in X-poisoning crosses and controls.

| gene/s targeted with sgRNA | nos-cas9 used | number of fathers used per vial | number of mothers per vial | Number of replicate vials used | Stock origin of mothers | number of sons | number of daughters | proportion of rarer sex | Drive efficiency (%)* |
| --- | --- | --- | --- | --- | --- | --- | --- | --- | --- |
| **None** | CAS-0001 | 1 | 3 | 14 | 09w | 1014 | 1081 | 0.48 | -3 |
| **None** | CAS-0001 | 1 | 1 | 14 | 09w | 603 | 544 | 0.47 | 5 |
| **None** | CAS-0001 | 1 | 3 | 4 | C(1)DX, y[1]f[1]/Y | 182 | 151 | 0.45 | 9 |
| **wupA** | CAS-0001 | 1 | 3 | 14 | 09w | 1409 | 176 | 0.11 | 78 |
| **wupA** | CAS-0001 | 1 | 1 | 14 | 09w | 698 | 94 | 0.12 | 76 |
| **wupA** | NH-noscas9 | 3 | 10 | 18 | 09w | 1569 | 257 | 0.14 | 72 |
| **wupA** | CAS-0001 | 1 | 3 | 6 | C(1)DX, y[1]f[1]/Y | 7 | 290 | 0.02 | 95 |
| **wupA** | NH-noscas9 | 3 | 10 | 18 | C(1)DX, y[1]f[1]/Y | 14 | 780 | 0.02 | 96 |
| **RpS19a+RpL35** | CAS-0001 | 1 | 3 | 14 | 09w | 789 | 195 | 0.20 | 60 |
| **RpS19a+RpL35** | CAS-0001 | 1 | 3 | 7 | C(1)DX, y[1]f[1]/Y | 55 | 143 | 0.28 | 44 |
| **RpL36+RpL35** | CAS-0001 | 1 | 3 | 14 | 10w | 678 | 717 | 0.49 | -3 |
| **RpL36+RpL35** | CAS-0001 | 1 | 3 | 6 | C(1)DX, y[1]f[1]/Y | 234 | 237 | 0.50 | 1 |

\*Two times the difference between the expected frequency of 50% and the observed % of rarer sex.

### *Wings-up A* poisoning skews the sex ratio

When the CRISPR machinery was targeted to *wupA,* significant skews in sex ratio towards maleness was observed (*p*<0.0001, t-test with Bonferroni correction, Figure 2b). However, we initially found the expression of sgRNAs targeting *wupA* did not result in a significant reduction in the total number of offspring produced when compared to controls (Fig. 2b). The highly significant reduction in the number of female offspring (*p*<0.0001, t-test with Bonferroni correction), was offset by the significantly higher number of male offspring compared to control (*p*<0.0001, t-test with Bonferroni correction). We hypothesised that the lack of reduction in reproductive output may be due to a carrying capacity within the vials with an excess of embryos being laid during the seven day period where remating was allowed. To address this possibility, single males heterozygous for both X-poisoning drive components (ie. the *wupA* targeting sgRNAs and nos-Cas9) were placed in vials with single females for four hours only. In these single pair crosses, a significant reduction in the total number of adult offspring (*p*<0.01, t-test with Bonferroni correction) as well as a highly significant change in sex ratio was observed (*p*<0.0001, t-test with Bonferroni correction; Figure 2c) and there was no significant difference in the number of males produced when compared to the control.

As *wupA* has very low expression in the early stages of embryos (first 8 hours; modENCODE Temporal Expression Data Flybase) it contrasts to many ribosomal proteins because it is not required for the viability of all cells and particularly not those in early development. The gene *wupA* encodes Troponin I, a protein that inhibits the interaction of myosin and actin in muscles although it is also thought to have non-muscle functions that relate to the nuclear localization of at least one of its nine isoforms (Casas-Tintó and Ferrús 2021, Flybase). To explicitly test where the sex-skewing of *wupA* CRISPR/nos-Cas9 elicited its effect we tracked the fate of X-chromosomes labelled with GFP. In the crosses we conducted, female embryos glowed green (after the 3XP3 GFP transgene is turned on), whereas male embryos and unfertilized eggs did not. Twenty single pair crosses were set up for the haplolethal cross (males bearing *wupA* sgRNAs and Cas9) and for controls (the males had *wupA* sgRNAs but lacked Cas9) and these revealed that the number of female embryos did not differ significantly between the haplolethal cross and the control cross, but the number of larvae, pupae and adults did (Figure 3). We could determine that females of the haplolethal cross typically died in late embryogenesis (dorsal closure was completed and we observed mouthparts and denticle belts however segmental furrows appeared unevenly spaced which may be attributable to abnormal muscle contraction; see Figure S1). Only ∼13% of GFP eggs became 1^st^ instar larvae, whereas ∼72% of GFP eggs from the control cross were observed to develop into larvae (Figure 3). Consistent with earlier results the ratio of females among adults in these experiments was ∼11% in the haplolethal cross and ∼52% in the control. We also examined 15 adult females emerging from this cross and all were fertile.

**Figure 3:**
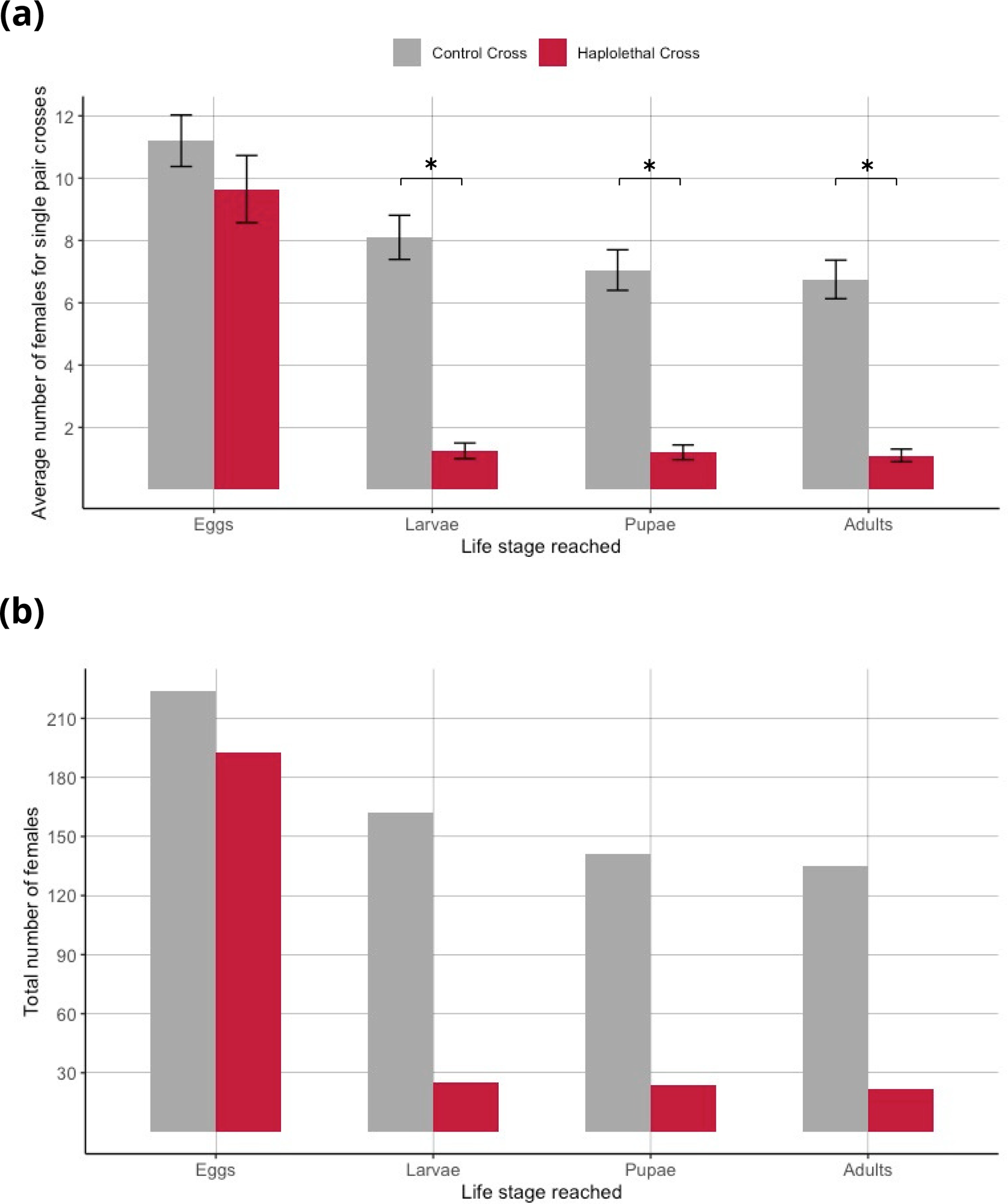
Developmental stage of lethality from *wupA* poisoned fathers. (a) The average number of daughters tracked from 20 vials of the haplolethal (*wupA*-poisoned) cross (in red) and 20 vials from a control cross (grey). Error bars represent the standard error of the mean and asterisks represent significance (P<0.05). (b) The total number of daughters from 20 vials from the haplolethal (red) and control cross (grey) at each life stage.

### No fitness cost detected in nos-Cas9 *wupA* poisoning males

Transcriptome datasets (e.g. FlyAtlas2-Anatomy RNA-Seq) indicate that *wupA* is expressed in the testis and so we also sought to determine whether CRISPR/Cas9 of *wupA* during spermatogenesis affected the fertility of the bearer. Males generating *wupA* alleles during spermatogenesis could have a phenotype that makes them less successful in mating, perhaps because of female choice, or they could produce lower quality sperm. To test for these possibilities we examined the fitness of males in a competitive assay. Four classes of test males (those with CRISPR/nos-Cas9 targeting *wupA*, and three control lines: one that carried the Cas9 without the sgRNA, another that had the sgRNA but no Cas9, and a third that had neither CRISPR component) individually competed for a female homozygous for the recessive ebony body colour mutation against an ebony male. Eighty-five vials containing two males and one female were set up (approximately 20 vials for each of the four tester male genotypes; Table 2) and the number of ebony and wildtype progeny counted. In the crosses involving males with both the sgRNA’s and Cas9 transgenes (cross A, Table 2) 84 of the 112 progeny were wildtype with ten of the former being females. So these test males were much more competitive than ebony males, even though there was a massive skew in the sex ratio so that only 12% of the progeny (10/84) were daughters (Table 2). In contrast, ebony males were more successful than the males of the three control crosses although only significantly so in cross D (Table 2).

**Table 2:**
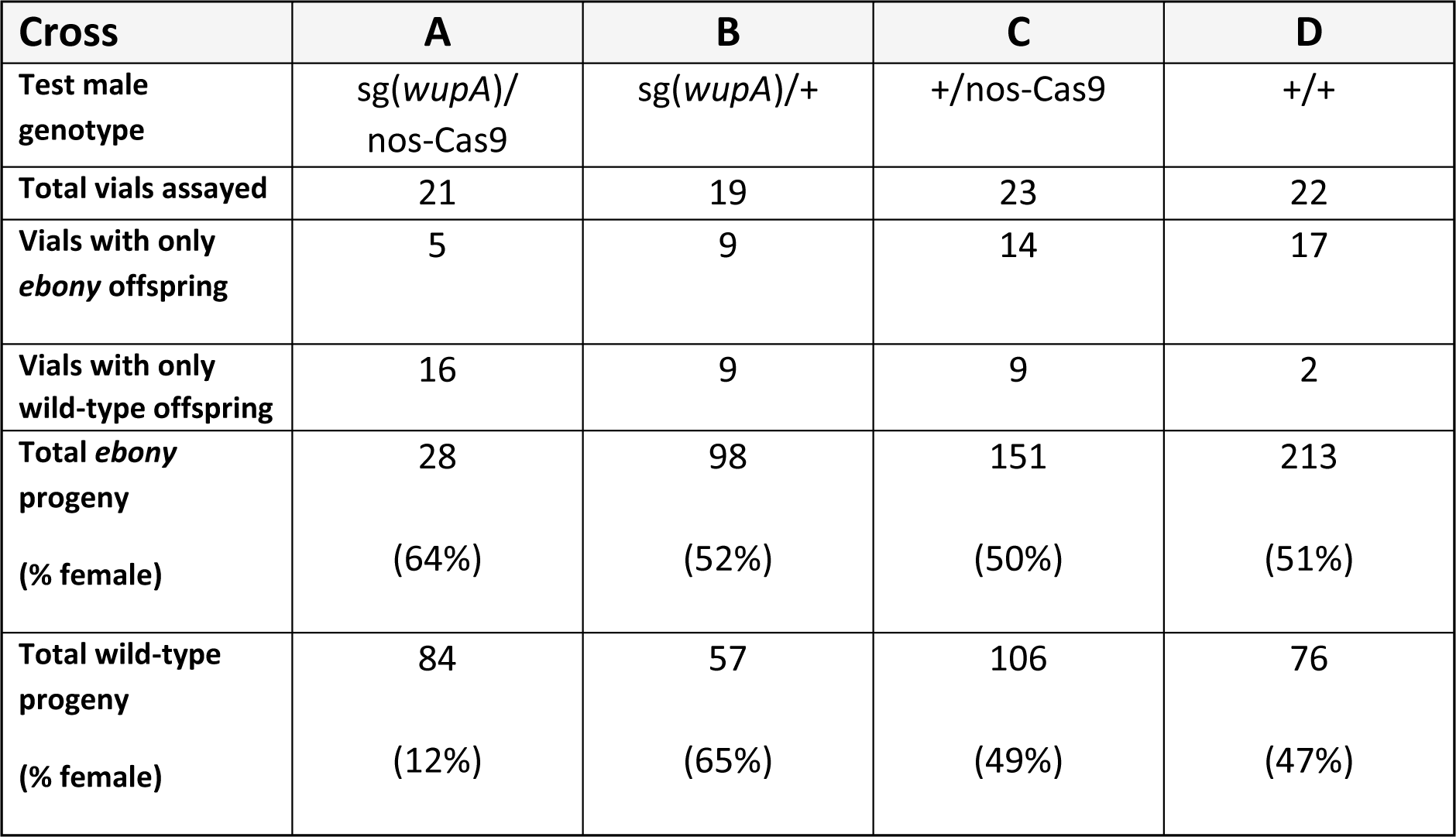
Competitive mating assays.

| Cross | A | B | C | D |
| --- | --- | --- | --- | --- |
| Test male genotype | *sg(wupA)/nos-Cas9* | *sg(wupA)/+* | *+/nos-Cas9* | *+/+* |
| Total vials assayed | 21 | 19 | 23 | 22 |
| Vials with only *ebony* offspring | 5 | 9 | 14 | 17 |
| Vials with only wild-type offspring | 16 | 9 | 9 | 2 |
| Total *ebony* progeny<br>(% female) | 28<br>(64%) | 98<br>(52%) | 151<br>(50%) | 213<br>(51%) |
| Total wild-type progeny<br>(% female) | 84<br>(12%) | 57<br>(65%) | 106<br>(49%) | 76<br>(47%) |

If competition is assessed at the level of vials then 81 of the 85 vials produce either ebony progeny or wildtype progeny. At this level the males with the full CRISPR/Cas9 machinery were also seen to be more successful than ebony males as they were the only sires in 16 vials, whereas ebony were the only sires in 5 of the vials (P=0.03). Again, this was in contrast to the crosses involving the three control genotypes where the number of ebony either did not differ significantly from wildtype (sgRNA-only control P=0.82; Cas9-only control P=0.21) or in the case of the background control line were more fit (P=7.6×10^-5^). So we found no evidence of a pre-laying fitness deficit for males that have CRISPR machinery targeting *wupA* during spermatogenesis in the *nanos* expression domain.

In contrast, we attempted to create *wupA*-poisoning males that would have Cas9 expressed under the *vasa* promoter, but this failed. We crossed males which had vasa-Cas9 on the Y chromosome and females that were homozygous for the *wupA* sgRNA construct. This yielded 264 daughters (that did not have Cas-9 because they did not inherit the Y chromosome) and only two small and infertile males. This is consistent with earlier reports that *vasa* expresses outside the germline (Gamez et al. 2021), and shows that X-poisoning with *wupA* could have drastic fitness costs if the CRISPR machinery is active in broader expression domains.

### The nos-Cas9 *wupA* sex biasing is not absolute

To understand why the sex biasing was not absolute, X-chromosomes that had survived being passed through the paternal germ line were characterized. If the targeted loci had been cut and reannealed, indels or single nucleotide mutations may have been introduced, and they would be revealed by sequence analysis at the sites targeted by the sgRNA’s. Initially, to simplify the sequence analysis, we used X^^^XY females (hereafter ‘attached-X’) which reverse the inheritance of the paternal X’s so that they are passed to sons. Consistent with expectations, males expressing *nanos*-Cas9 and sgRNA’s targeting *wupA* sired significantly more daughters than sons (the sex ratio bias being flipped due to sons paternally inheriting the X chromosome; *p*<0.001, t-test with Bonferroni correction). The RpS19a+RpL35 cross showed no significant sex ratio bias but had significantly fewer male offspring than the control and no significant difference in the number of females (Figure 2d; *p*<0.0001, t-test with Bonferroni correction). There was no reduction in the number of offspring produced by males expressing gRNA targeting RpL36+RpL35.

The X chromosomes of a total of 60 surviving sons (14 for *wupA*, 22 for RpS19a+RpL35 and 24 for RpL36+RpL35) from 15 independent crosses were Sanger sequenced and only the offspring of one male showed any sign of CRISPR cutting. A complex indel resulting in a net 9 nucleotide insertion that did not introduce a premature stop codon was recovered from a cross involving the RpL36+RpL35 sgRNAs (Figure S2). The fact that this individual survived suggests that gene drive resistance alleles could arise against RpL35 gRNA1. The data also concurs with the idea that these genes are indeed lethal genes as no frameshifting indels or nonsense mutations were recovered. More generally, these data are consistent with the notion that most of the surviving progeny carrying a paternal X never had their X chromosome cut by the CRISPR machinery.

### Not all *wupA* mutations are haplolethal

Curiously, we found that the skew in sex ratio was more extreme when the *wupA*-poisoned X chromosome was passed to sons (2%; via attached-X females) rather than to daughters (14%; Figure 2d versus Figures 2b and 2c; Table 1). As the fathers in these crosses are the same, this difference is not due to the generation of X-poisoned alleles. Instead, heterozygosity in daughters may protect against lethality more than the exposed hemizygosity of sons, which suggests that some of the CRISPR generated alleles may encode recessive lethals. If true then we reasoned it might be possible to find classes of mutations in daughters of males that have *wupA* targeted during spermatogenesis that do not occur in their sons (from crosses with attached-X). Three PCR amplicons surrounding the three sgRNA sites in *wupA* were sequenced from twenty pools of ten daughters using Illumina’s short read MiSeq sequencing technology. Strikingly, each of the 20 pools had a high frequency of deletion alleles (between 5 and 18%) among the reads from the amplicon that amplified across the site targeted by sgRNA2 (Figure 4). Given that each of the 20 pools was composed of ten daughters, and we saw no such indels among the 14 sons from the attached-X cross, a significant number of the mutations generated by CRISPR-editing seem to be lethal (purged in sons) but not haplolethal (and so present in sequences from daughters). Notably all of the high frequency indels observed had a length divisible by three and therefore did not change the open reading frame. So it would seem that frameshift alleles are haplolethal but some indels that maintain the open reading frame are lethal but not haplolethal.

**Figure 4:**
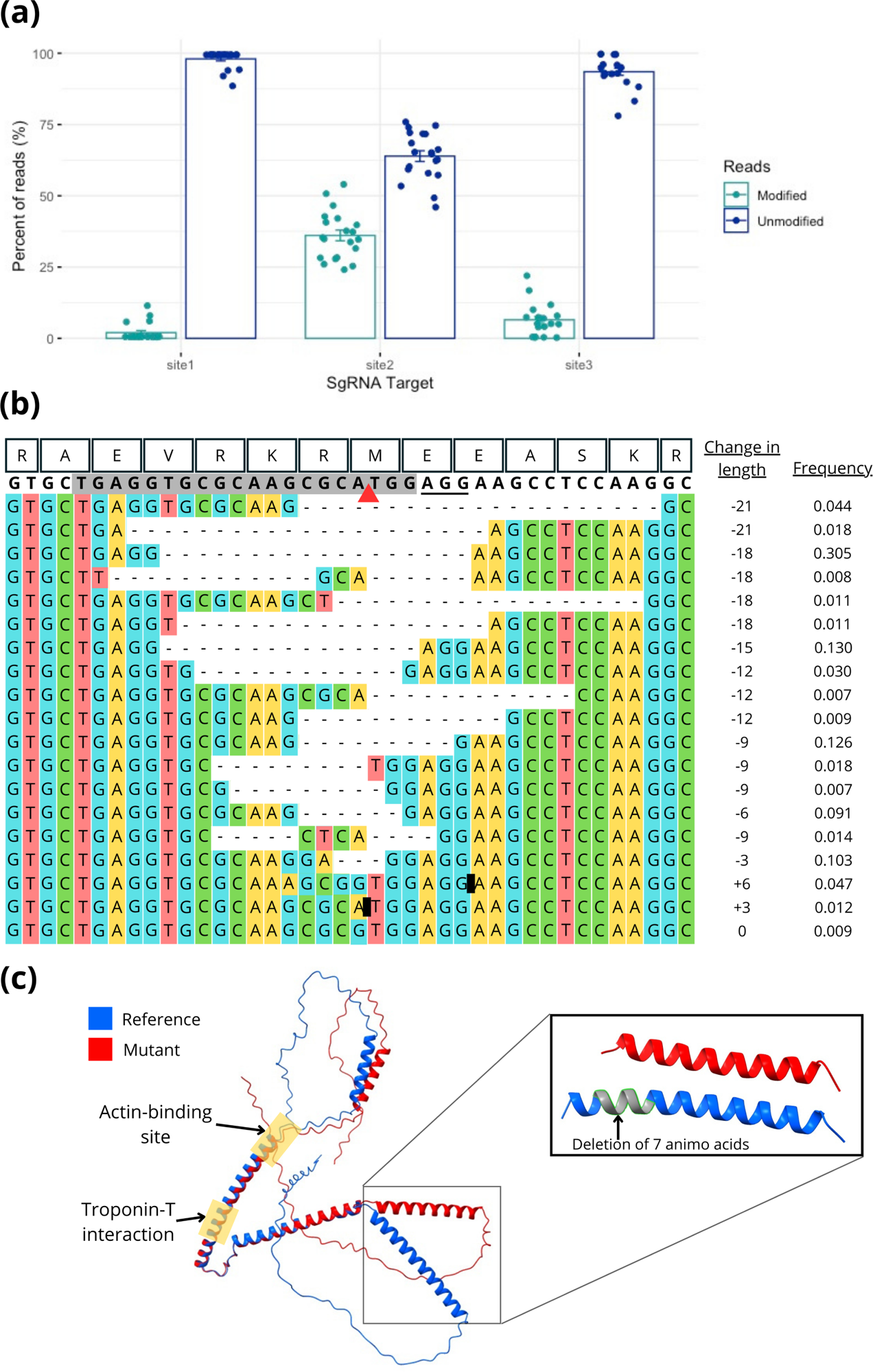
Recessive lethal *wupA* mutations revealed in females. (a) The three sites within *wupA* that are targeted by sgRNAs (see figure 1c) vary in the number of CRISPR mutations uncovered by sequencing 200 daughters of an X-poisoning cross. Here a Crispresso2 output shows the total number of modified and unmodified reads from Illumina sequencing at each target site. (b) A nucleotide alignment of the modified reads of target site 2 showing their frequencies. Amino acid sequences are shown at the top, the sgRNA sequence is highlighted in grey, the PAM site is underlined, the red arrow is the cut site, and black blocks represent insertions (shown more to detail in Figure S3). (c) Isoform G of Tropinin I without any mutations (blue) and with the largest deletion found in our sequencing results (a 21 bp deletion shown in red). Protein folding predicted with AlphaFold. The deletion location is highlighted in grey. No mutations in target site 2 will affect the acting-binding and troponin-interacting areas.

## Discussion

For an X-poisoning strategy to elicit an effect in a single generation the targeted genes need to be haplo-insufficient. Such is the knowledge associated with *Drosophila melanogaster* that it is known there are eleven loci on the X chromosome of *D. melanogaster* that are haplolethal or haplosterile (Cook *et al*. 2012). Haplolethal genes can only exist on the X chromosome in species such as Diptera where dosage compensation between the homogametic and heterogametic sex is due to upregulation - doubling the output of the X chromosome in males, rather than the inactivation of the X as seen in humans.

Eight of the X-linked haplolethals of *D. melanogaster* encode ribosomal proteins, which contribute to the translational machinery of cells, and an insufficient quantity of them leads to lethality or to the Minute syndrome where flies are tiny and sterile (Marygold *et al*. 2007). Previously, Fasulo *et al*. 2020 pioneered the ‘X-poisoning’ strategy by targeting Ribosomal protein genes S6 and S5a. They found that when one particular sgRNA targeting RpS6 was combined with Cas9 driven by a sperm-specific beta tubulin 85D promoter the sex ratio was biased so that greater than 92% of progeny were males. Our targeting of RpL36, RpL35 and RpL19a did not produce such extreme biasing. However, Fasulo *et al*. (2020) also showed heterogeneity in the success of their sgRNA’s with different guide RNA’s targeting the same genes skewing to various extents. Therefore, the failure to see skewing in our combination of RpL36 and RpL35 sgRNA’s or the skew to only 80% males in the RpL35 and RpL19a combination, may well have more to do with the particularities of guide RNA efficacy than being a property of the genes they target, and in our case by subtle design features such as the position of particular guide RNAs in the array may be important.

A surprising finding of Fasulo *et al*. (2020) was that some alleles among the survivors contained frameshift mutations. This led the authors to suggest that the targeted ribosomal protein genes may not be haplolethal genes after all and that previously studied alleles may have harboured dominant negative mutations instead. In our analyses, we did not find such frameshifting alleles among individuals carrying X-chromosomes that had passed through X-poisoning males. Instead, the loci showed no signs of being cut at all or exhibited mutations that maintained the frame (Figure 4b for sgRNA2 of *wupA* and Figure S3 for sgRNA1 and sgRNA3 of *wupA*).

The *wupA* locus of Drosophila was the only non-ribosomal protein haplolethal gene that has been molecularly characterised on the X chromosome and it showed the greatest skewing in our X-poisoning experiments. It encodes Troponin I, a protein that inhibits the interaction of myosin and actin in muscles. It is also thought to have non-muscle functions that relate to the nuclear localization of at least one of its isoforms (Casas-Tintó and Ferrús 2021, Flybase). The gene has 12 exons that are variously combined to produce 13 alternate transcriptional splice forms. Two of the three sgRNAs used in our study target exons common to all splice forms (sgRNA2 and sgRNA3), and the third targets a PAM site that is found in the first two codons of isoforms with the most common start methionine. The latter site was previously targeted with CRISPR by Casas-Tintó and Ferrús (2021) and the resulting haplolethality is consistent with our findings.

Our results provide an insight into the distinct functions of *wupA* because indels were enriched around the sgRNA2 target site which lies on top of the methionine codon used by *wupA* isoforms that translocate into the nucleus. We infer that such mutations are recessive lethals that are exposed in the hemizygous condition of males. However, as heterozygotes in females they produce Troponin I with an intact actin-binding site and an intact Troponin T interaction site and so they are not haplolethal. The haplolethality of *wupA* seems to relate to the delicate stoichometry required for Troponin I and F-actin. Without sufficient Troponin I, calcium is not required to allow tropomyosin to move and allow actin to interact with myosin. Thus, lethality arises from the irreversible hyper-contraction of muscles. Such a model is consistent with observations that mild *wupA* alleles (yielding the wings-up phenotype) can be rescued by a mild allele of tropomyosin (Naimi et al. 2001).

There is much about *wupA* that makes it an attractive target for future X-poisoning strategies. As a haplolethal gene not expressed until late embryogenesis, the progeny of a poisoned cross are unaffected at fertilization and egg-lay. It thus contrasts with X-poisoning targeting ribosomal proteins which may have reduced fertility because spermatogenesis may be affected. It also contrast with genetic systems causing sterility (such as PG-SIT (Kandul et al. 2019)) which can be counteracted by females mating multiple times. We also did not find males that poison *wupA* to have reduced fitness, although we acknowledge that our competitive assays are likely be confounded with genetic background effects (especially eye-colour attributable to the number of transgenes) but argue that if there is any fitness cost attributable to *wupA* poisoning using a *nanos*-driven Cas9 it is less than the loss of a second transgenic copy of *mini-white* and less than that of ebony males. The *wupA gene* is a highly conserved gene that is found on the X chromosome on numerous dipteran species, including some important pests (Figure S4). *Drosophila suzukii,* an invasive pest of soft-skinned fruits, appears to share all 12 exons with *D. melanogaster* and all the predicted proteins have greater than 98% amino acid identity with *D. melanogaster* homologs. In fact, two of the three sgRNA’s used here in *D. melanogaster* match *D. suzukii* targets perfectly, and the third has only one nucleotide different. *wupA* of Anopheles mosquito’s has a slightly different gene structure (eg. Exon 1 and Exon 2 are fused eg. XM_053811545) however it is also found on the X chromosome in these species suggesting it could be a target of X-poisoning strategies in the many pest species of this genus. Unfortunately, *wupA* orthologs are not on the X-chromosome of all economically important dipterans, such as the tephritids which have Muller element F (chromosome 4 of Drosophila) as an X chromosome.

Taken together the above results suggest that *wupA* could be a good target for an X-poisoning gene drive. To create such a drive, transgenes encoding a germline restricted Cas9 and the sgRNA’s targeting *wupA* would ideally be placed on the Y chromosome. Placing transgenes on the Y chromosome has been technically challenging although it has been done (Gamez *et al*. 2021). Indeed, we examined the one *D. melanogaster* line currently available that has a Cas9 transgene on the Y chromosome however because it uses a *vasa* promoter that expresses outside the germline males that poisoned the X during spermatogenesis could not be generated. An important next step in this research program will be to overcome the challenge of placing germline-limited drive components onto the Y chromosome.

Finally, we note that where an X-poisoning strategy can take the form of a Y-linked editor (Burt and Deredec 2018) it may not necessarily drive through a population. While fathers may only have sons, the Y-chromosome bearing the X-poison may not produce more sons than any other Y-chromosome in a population and so it may not be considered a gene drive. However, if X-poisoned males produce more sons than other Y-chromosomes, perhaps as a result of reduced larval competition with sisters (such as seen in the original *wupA* cross), then a Y-linked editor could indeed be a gene drive. In this larval-competition scenario, as the frequency of the Y-linked editor increases, then so too will the strength of the gene drive.

## Data Availability Statement

All data is here or in the supplementary material.

## Acknowledgements

We’d like to thank Owain Edwards, Max Scott and Nina Wedell for conversations that helped develop this research and Jackson Champer for suggestions about construct design. We also thank the anonymous reviewers for suggestions that improved the manuscript.

